# A comprehensive ensemble model for comparing the allosteric effect of ordered and disordered proteins

**DOI:** 10.1101/377135

**Authors:** Luhao Zhang, Maodong Li, Zhirong Liu

## Abstract

Intrinsically disordered proteins/regions (IDPs/IDRs) are prevalent in allosteric regulation. It was previously thought that intrinsic disorder is favorable for maximizing the allosteric coupling. Here, we propose a comprehensive ensemble model to compare the roles of both order-order transition and order-disorder transition in allosteric effect. It is revealed that the MWC pathway (order-order transition) has a higher probability than the EAM pathway (disorder-order transition) in allostery, suggesting a complicated role of IDPs/IDRs in regulatory proteins. In addition, an analytic formula for the maximal allosteric coupling response is obtained, which shows that too stable or too unstable state is unfavorable to endow allostery, and is thus helpful for rational design of allosteric drugs.

**Author Summary:** Allosteric effect is an important regulation mechanism in biological processes, where the binding of a ligand at one site of a protein influences the function of a distinct site. Conventionally, allostery was thought to originate from structural transition. However, in recent years, intrinsically disordered proteins (IDPs) were found to be widely involved in allosteric regulation in despite of their lack of ordered structure under physiological condition. It is still a mystery why IDPs are prevalent in allosteric proteins and how they differ from ordered proteins in allostery. Here, we propose a comprehensive ensemble model which includes both ordered and disordered states of a two-domain protein, and investigate the role of various state combinations in allosteric effect. By sampling the parameter space, we conclude that disordered proteins are less competitive than ordered proteins in performing allostery from a thermodynamic point of view. The prevalence of IDPs in allosteric regulation is likely determined by all their advantage, but not only by their capacity in endowing allostery.

## Introduction

Allosteric regulation is intrinsic to the control of many metabolic and signal-transduction pathways.(1) It is described as the effect that the binding of a ligand at one site of a protein influences the function of a distinct site which binds with substrate.(2) In history, several models have been proposed illuminating possible mechanism of allostery. The classical MWC (Monod-Wyman-Changeux)(3) model explained the allosteric effect based on a cooperative conformational transition of protein oligomers. Taking hemoglobin binding with oxygen as an example [see Fig. 1(a)], the MWC model assumes that four subunits of hemoglobin are simultaneously in either a relaxed state (R state) or a tense state (T state), and oxygens bind preferentially to the R state which shifts the R-T equilibrium. With such a simple assumption, the MWC model nicely explained how the binding of oxygen at one site promotes the binding at a remote site. Later, the KNF (Koshland-Nemethy-Filmer) model(4) has considered finite subunit interactions and proposed a progressive conformational transition of each domain step by step [Fig. 1(b)]. Both models imply that allosteric processes are closely associated with ligand-driving conformational changes that propagate between the allosterically coupled binding sites. With the development of structural biology, the description of allostery in terms of structure changes was derived,(5) and was used to study allosteric proteins such as lactate dehydrogenase.(6) The structure paradigm also leads to the seeking of specific atomic pathway that connects allosteric sites.(7, 8) Nevertheless, the discovery of dynamic structure and multiple conformations of proteins, such as multiple orientations of DNA-binding domains of DNA-binding proteins in the absence of DNA(9) and the intermediate conformation of hemoglobin in solution,(10) suggests more possibilities beyond the simple two-state models.

**Fig 1.**
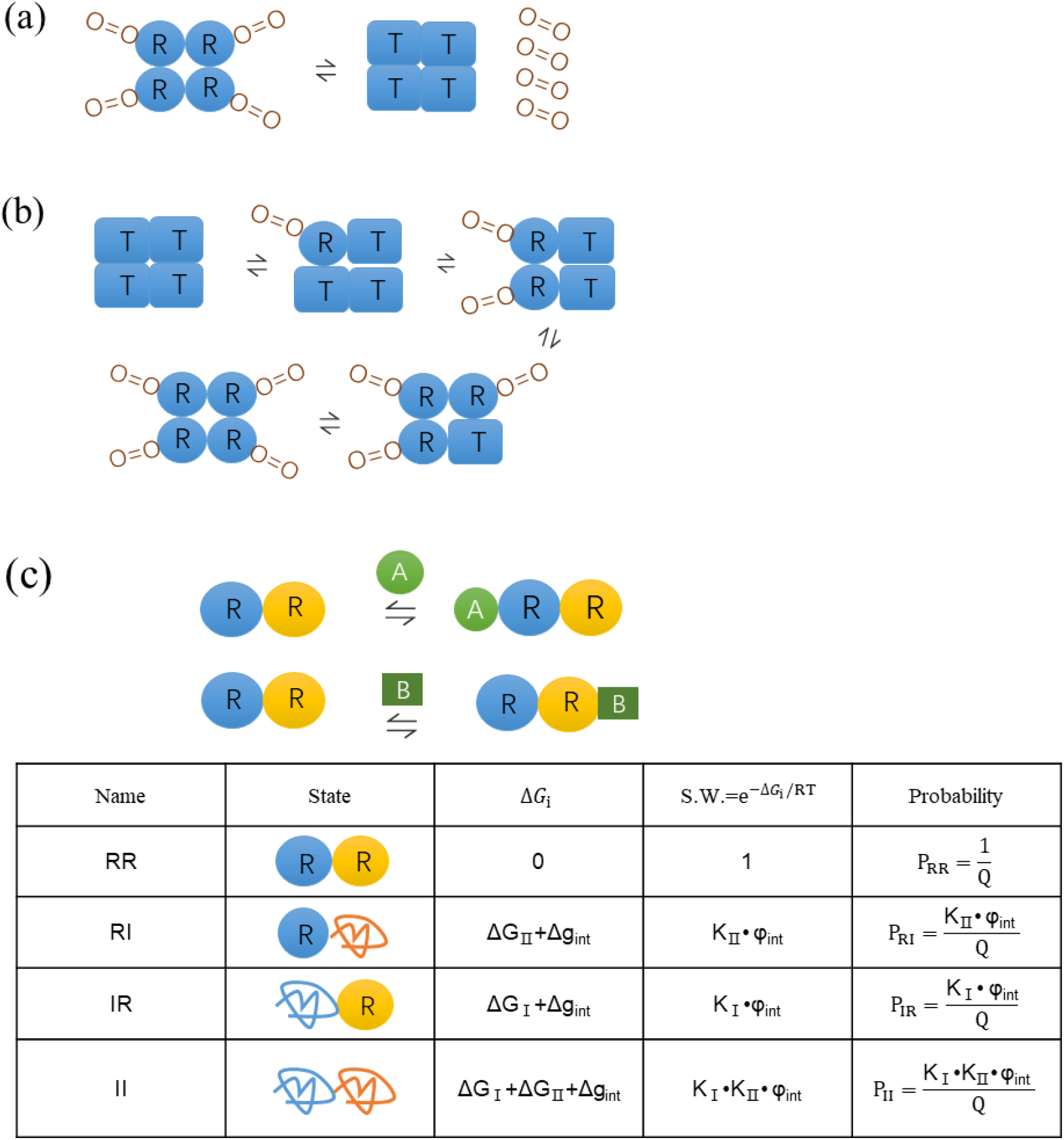
Schematic diagram of MWC, KNF and EAM models. (a) The MWC model for hemoglobin.(3) All four subunits are simultaneously in either a R (Relaxed) state (with a higher affinity for O_2_) or a T (Tense) state (with a lower affinity for O_2_). The cooperative effect happens when one O_2_ binds to the all-R state, shifts the chemical balance from all-T to all-R, then creates more favorable binding sites for subsequent O_2_. (b) The KNF model for hemoglobin.(4) It allows several intermediate states keeping balanced by several chemical equilibrium constants. (c) The EAM model for a two-domain protein.(11) Each domain can be in either a R (Relaxed) state or an I (Disordered) state, resulting in four possible combinations for protein states. The free energy Δ*G_i_* (relative to the RR state), the statistical weight (S.W.) and the corresponding probability of four states were listed. One domain (blue) in the R state can bind the allosteric ligand (A) while the other domain (yellow) in the R state can bind the substrate (B). The I state of each domain has no affinity to ligand and substrate.

The discovery of intrinsically disordered proteins (IDPs) and intrinsically disordered regions (IDRs) has brought a challenge to the conventional “structure-function” paradigm.(12–15) IDPs/IDRs do not have ordered structures in the free state under physiological conditions, but they are important in biological signaling and regulation.(16–23) IDPs/IDRs possess some advantages over ordered proteins,(24) such as high specificity coupled with low affinity useful for reversible signaling interaction,(25–28) binding to multiple partners,(29, 30) and rapid turnover allowing sensitive response to environment changing.(12, 19, 31) Therefore, they play crucial roles in widespread categories of proteins,(22) e.g., scaffold proteins,(32) RNA and protein chaperones,(33) transcription factors,(20) and regulation of cellular pathways.(34) In particular, IDPs/IDRs were found to be widely involved in allosteric regulation in despite of their lack of ordered structures.(35–42) Representative examples include enzyme aminoglycoside N-(6’)-acetyltransferase II (AAC), which has local unfolding and switching behaviors from positive cooperativity to negative cooperativity upon different temperature;(37) and Doc/Phd toxin-antitoxin system with intrinsic disorder exhibiting complex “conditional cooperativity” character upon different Doc/Phd ratio.(38)(42)

How can IDPs/IDRs implement allosteric effect under the lack of ordered structures? And why are they so prevalent in allosteric regulation? The answer is related to an emerging new view of allostery based on the general landscape theory of protein structure, where the ligand binding stabilizes specific states and shifts the conformational ensemble.(11, 43, 44) The EAM (Ensemble Allostery Model) model used the ensemble view to explain the allostery of IDPs,(45–49) see Fig. 1(c). As an example, it described a two domain system as a four-state ensemble with each domain having ordered (R) and disordered (I) states. The allosteric ligand (A) binds only with the R state of the first domain while the substrate (B) binds only with the R state of the second domain, i.e., the disordered states have no affinity to ligand and substrate. When the interface-interaction free energy between two ordered domains is negative, binding of the ligand A would stabilize the RR state and thus facilitate the binding of the substrate B, resulting in a positive allosteric effect. Similarly, a negative allosteric effect arises when the interface interaction is unfavorable. The EAM model also provided insight in explaining why IDPs/IDRs are so prevalent in allosteric regulation: it was shown that high allosteric intensity is accompanied by high probability of disordered (I) states.(45) However, based on the ensemble concept, EAM model considers only the order-disorder (R-I) transition, but lacks the order-order (R-T) transition as that in the MWC model for the allostery of ordered proteins. Therefore, with separate EAM or MWC models, it is impossible to determine whether disordered or ordered proteins are more advantageous in allosteric regulation. To get a full view of competition of ordered and disordered proteins in allosteric effect, here we propose a comprehensive ensemble model considering both order-disorder and order-order transitions. In this comprehensive model, the EAM and MWC mechanisms become two pathways for allostery of the system, and thus their role can be quantitatively evaluated.

## Models

### The comprehensive ensemble model

Our proposed model describes a two-domain protein system, see Fig. 2. It combines components of both the MWC and the EAM models. Each domain has three states: R (Relaxed), T (Tense) and I (Disordered). Being consistent with the MWC model, R and T are incompatible and thus the combinations “RT” and “TR” are forbidden in the resulting protein states. Similar to the EAM model, the I state of a domain is disordered and do not have any interface interaction with the adjacent domain, and it does not bind to any ligand or substrate due to the lack of ordered structures. As a result, there are seven possible combinations for protein states, which are listed in Fig. 2 with the formula of their free energy, the statistical weight and the corresponding probability in the absence of ligand and substrate. Six free energy parameters (Δ*G*_R1_, Δ*G*_R2_, Δ*g*_int,R_, Δ*g*_int,T_, Δ*G*_RT1_, Δ*G*_RT2_) are basic parameters of the model, determining the ensemble distribution. The substrate B binds only to the R state of one (yellow) domain. The allosteric ligand A binds to the other (blue) domain but there are two binding modes: in the A-R binding mode A binds only to the R state of the blue domain, while it is the A-T binding mode when A binds only to the R state of the blue domain. The two binding modes are taken into account here to enable both positive and negative allosteric effects for ordered proteins (MWC mechanism), making a comparison between the roles of ordered and disordered proteins possible. For example, if we look at a subsystem consisting of RR and TT states, binding of A in the A-R binding mode increases the fraction of the RR state and thus enhances the subsequent binding of B (activation), while that in the A-T binding mode weakens the binding of B (inhibition).

**Fig 2.**
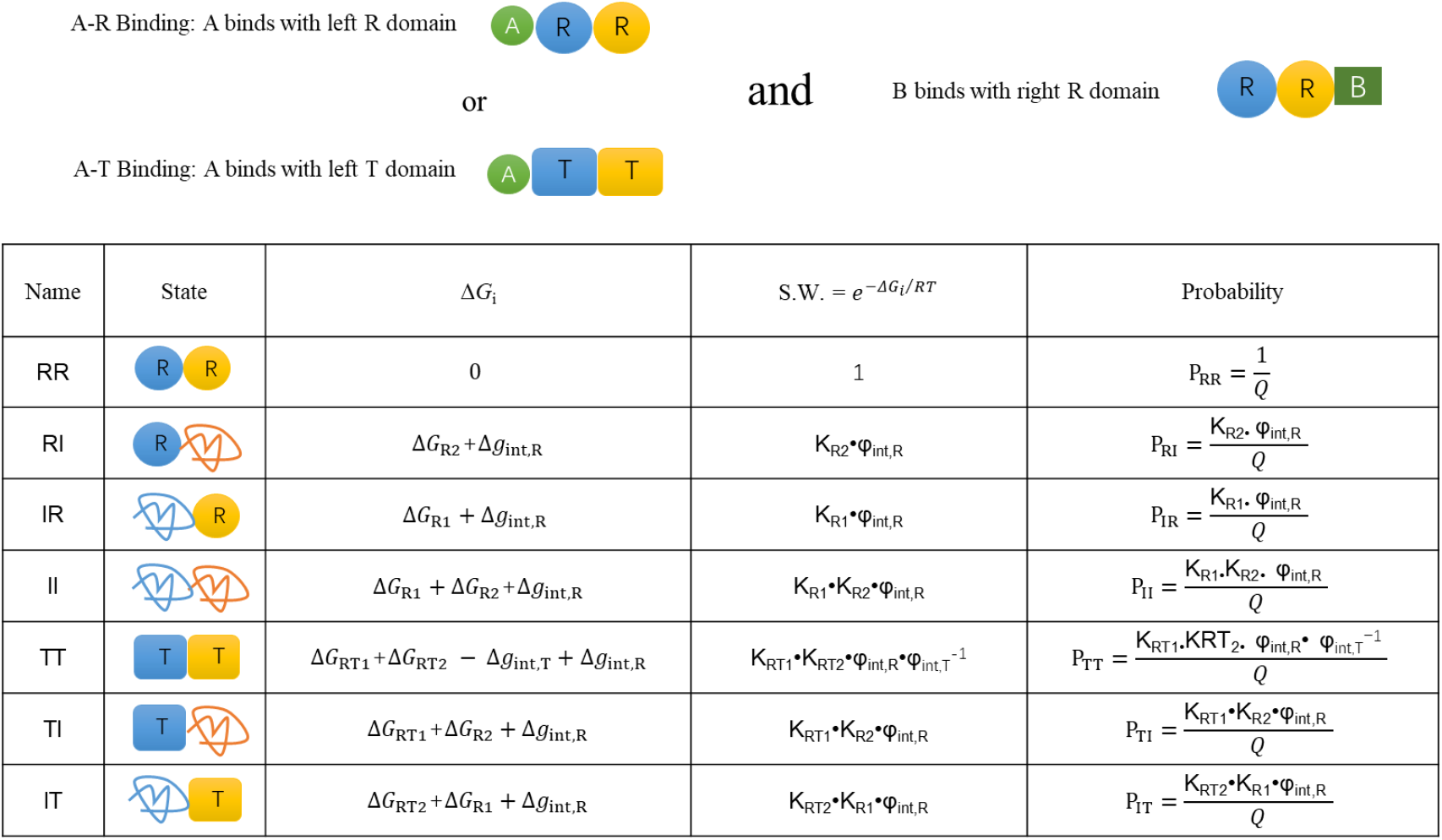
Schematic representation of the proposed comprehensive ensemble model. Each domain (blue and yellow) can be in R (Relaxed), T (Tensed) and I (Disordered) state. R and T are incompatible and thus there are seven possible combinations for protein states. They were listed with the free energy (relative to RR as the reference state as that in the EAM model), the statistical weight (S.W.) and the corresponding probability. Δ*G*_R1_ and Δ*G*_R2_ are the free energy of unfolding the R state of each domain, and Δg_int,R_ and Δg_int,T_ are the free energy of breaking the interface interactions in RR and TT, respectively, which were defined in a manner similar to the EAM model. Δ*G*_RT1_ and Δ*G*_RT2_ are the free-energy of the R-T transition for each domain. The statistical weights are given with 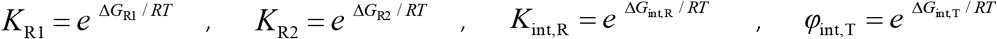, 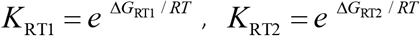. Q is the sum of statistical weights of all the states: 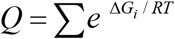. A is allosteric regulation ligand binding to one (blue) domain, and B is the substrate to the other (yellow) domain. A and B are different molecules, i.e., we consider the heterotropic allosteric effect. To enable both positive and negative allosteric effect for ordered proteins, we consider two binding modes for A: it can only bind to the R state of the blue domain (A-R binding mode), or can only bind to the T state of the blue domain (A-T binding mode). B always binds only to the R state of the yellow domain.

### Definitions of contribution of ordered and disordered protein pathways to allostery of the comprehensive ensemble model

Adding ligand A to the system results in a redistribution of the protein ensemble probabilities. The allosteric effect is directly related to probability variation of the states that can bind substrate B due to the adding of A. Following the EAM model,(45) we define the allosteric coupling response (*CR*) as

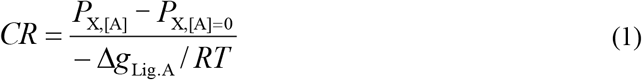

to quantitatively measure the allosteric intensity for a given system. Here, X denotes the states that can bind B, so *P*_X,[A]_ is the probability of states that can bind B when there exists ligand A, and *P*_X,[A]=0_ is the probability when A is absent. In the comprehensive ensemble model proposed here, for the A-R binding mode we have *P*_X,[A]_ = *P*_ARR_ + *P*_RR_ + *P*_IR_, and for the A-T binding mode we have *P*_X,[A]_ = *P*_RR_ + *P*_IR_. Δ*g*_Lig.A_ is the stabilizing free energy of adding ligand A for the states that can bind A, which is determined as:

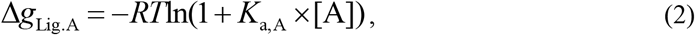

where *K*_a,A_ is the intrinsic equilibrium constant of the binding reaction for A. For example, in the A-R binding mode, *K*_a,A_ is the association constant for the reactions A + RR = ARR and A + RI = ARI, which gives the equilibrium distributions:

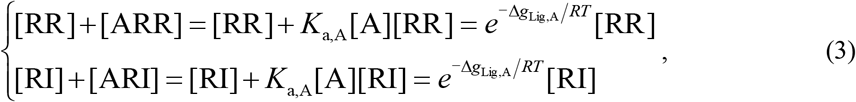

clearly demonstrating the nature of the stabilizing free energy Δ*g*_Lig,A_. In our study, we fixed Δ_Lig.A*g*_= −3.0 kcal/mol at a physiological temperature of *T* = 310.15 K unless otherwise specified.

Because the comprehensive model includes all the states of the MWC model and the EAM model, we can also view the comprehensive system consisting of three subsystems: the MWC subsystem, the EAM subsystem and the Others subsystem, and thus the allosteric effect can be approximately decomposed into three pathways (Fig. 3). The MWC pathway is order-order transition involving the states RR and TT, the EAM pathway is disorder-order transition involving the states RR, RI, IR and II, and the Others pathway is an extra component in the comprehensive model involving RR and the remaining states (TI and IT) neglected in the MWC and EAM pathways. The allosteric coupling response (CR) of each subsystem can be defined and calculated separately. Take the MWC subsystem as an example (under A-R binding mode), we have

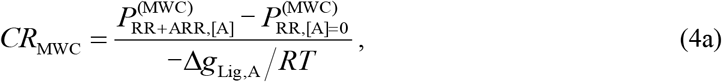

where the superscript “(MWC)” indicates that the related probabilities of states are defined (normalized) within the MWC subsystem. Similarly, *CR* for the EAM subsystem and the Other subsystem are determined by

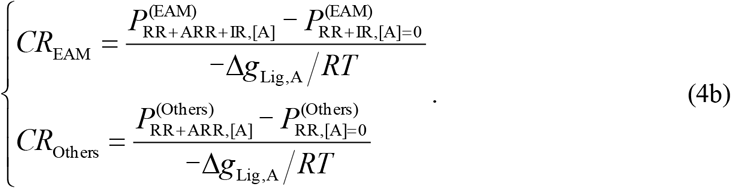

**Fig 3.**
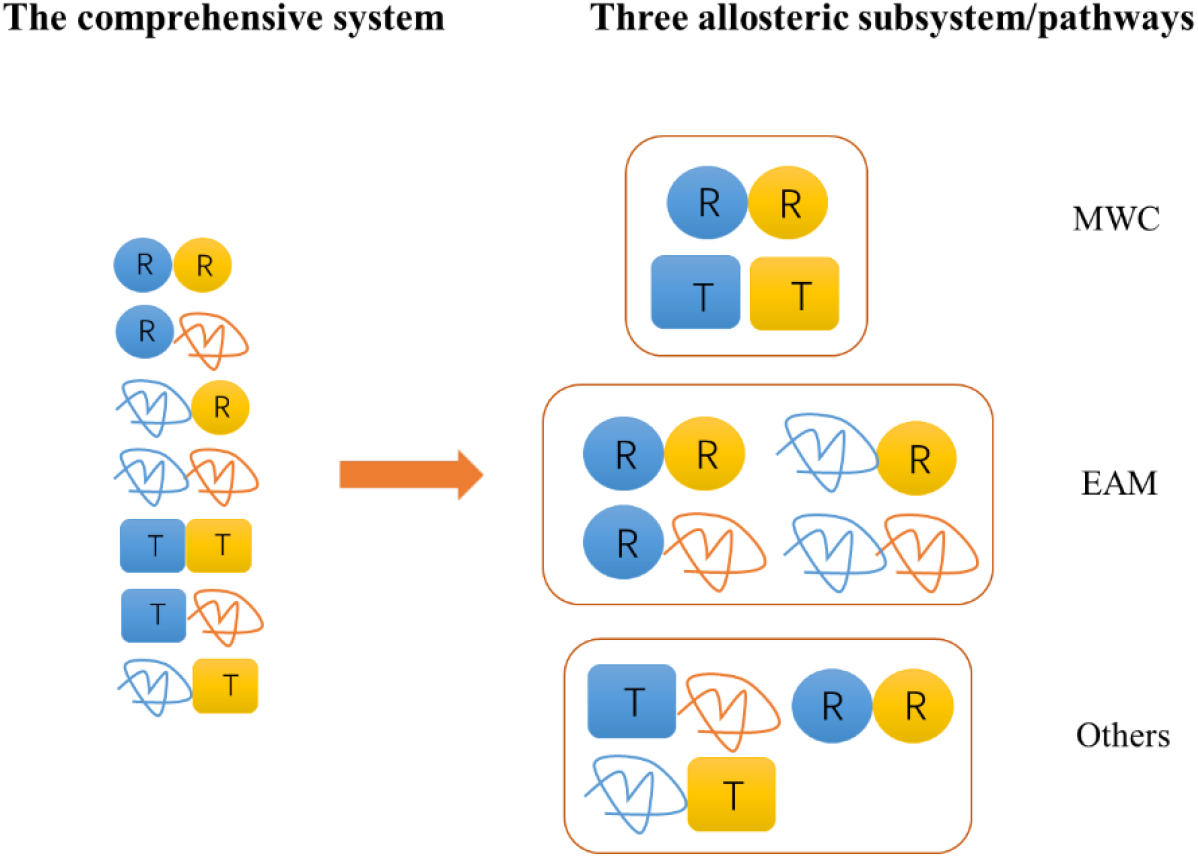
Three allosteric subsystems/pathways in the comprehensive ensemble model. The contribution ratios of pathways to the allostery of the comprehensive system are given in Eq. (5).

With a set values of the basic parameters (Δ*G*_R1_, Δ*G*_R2_, Δ*g*_int,R_, Δ*g*_int,T_, Δ*G*_RT1_, Δ*G*_RT2_), it is thus straightforward to calculate the probabilities of all the states with and without ligand A, as well as *CR* for the whole system (*CR*_tot_) and subsystems (*CR*_MWC_, *CR*_EAM_, *CR*_Others_). The contribution of a pathway to the total allostery of the comprehensive system depends not only on *CR* of the corresponding subsystem, but also on the proportion of the subsystem states in the whole system. Therefore, the contribution ratio of the MWC pathway to the allostery of the comprehensive system is approximately defined as:

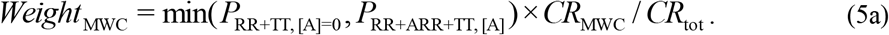

It stands for the weight of the MWC pathway in the allosteric effect. When there are only RR and TT states before adding ligand A, the comprehensive model degenerates to the MWC model and Eq. (5a) gives *Weight_MWC_* = 1. Similarly, for the EAM and the Others pathways, we have:

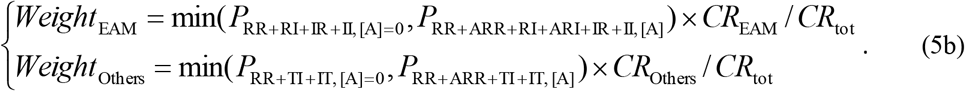

It is noted that *Weight*_MWC_, *Weight*_EAM_ and *Weight*_Others_ are metrics for three pathways’ contributions to allosteric effect of the comprehensive system, but the sum of them is not necessarily equal to 1.0 although the deviation is usually small. Related equations under the A-T binding mode can be found in Supporting Information.

## Results

### Limits for the maximal allosteric response

With a given set of parameters for protein state stability (Δ*G*_R1_, Δ*G*_R2_, Δ*G*_RT1_, Δ*G*_RT2_, Δ*g*_int,R_, Δ*g*_int,T_) and protein-ligand interaction (Δ*g*_Lig,A_) of the proposed comprehensive ensemble model, we can calculate the ensemble distribution, the allosteric coupling response (*CR*) and the contributions of different pathways with the formulism described above. *CR* as a function of Δ*g*_int,R_ and Δ*g*_int,T_ is shown in Fig. 4(a,b) as a case example when the other parameters are fixed. It reveals that combination of Δ*g*_int,R_ and Δ*g*_int,T_ is required to maximize the allosteric effect. Under the A-R binding mode, the model can afford both positive (*CR* > 0) and negative (*CR* < 0) allosteric effects, while there is only negative effect under the A-T binding mode. The achieved highest *CR* is about 0.17. To have a global inspection on the occurring probability of allostery, we assume the stability free-energy parameters (Δ*G*_R1_, Δ*G*_R2_, Δ*G*_RT1_, Δ*G*_RT2_, Δ*g*_int,R_, Δ*g*_int,T_) vary randomly between [−8, +8] kcal/mol, and determine the distribution of *CR* for two binding modes with Δ*g*_Lig,A_ = −3 kJ/mol [Fig. 4(c,d)]. For the majority of parameter sets, the resulting allostery is weak, giving a sharp peak at *CR* = 0 for both binging modes [Fig. 4(c,d)]. Actually, only 6.3% of parameter sets produce |*CR*| >0.1 under the A-R binding mode. Remarkably, *CR* has the boundaries at around ±0.172. In other words, no matter how the state stabilities of protein are optimized, it is impossible to achieve a *CR* value higher than 0.172.

**Fig 4.**
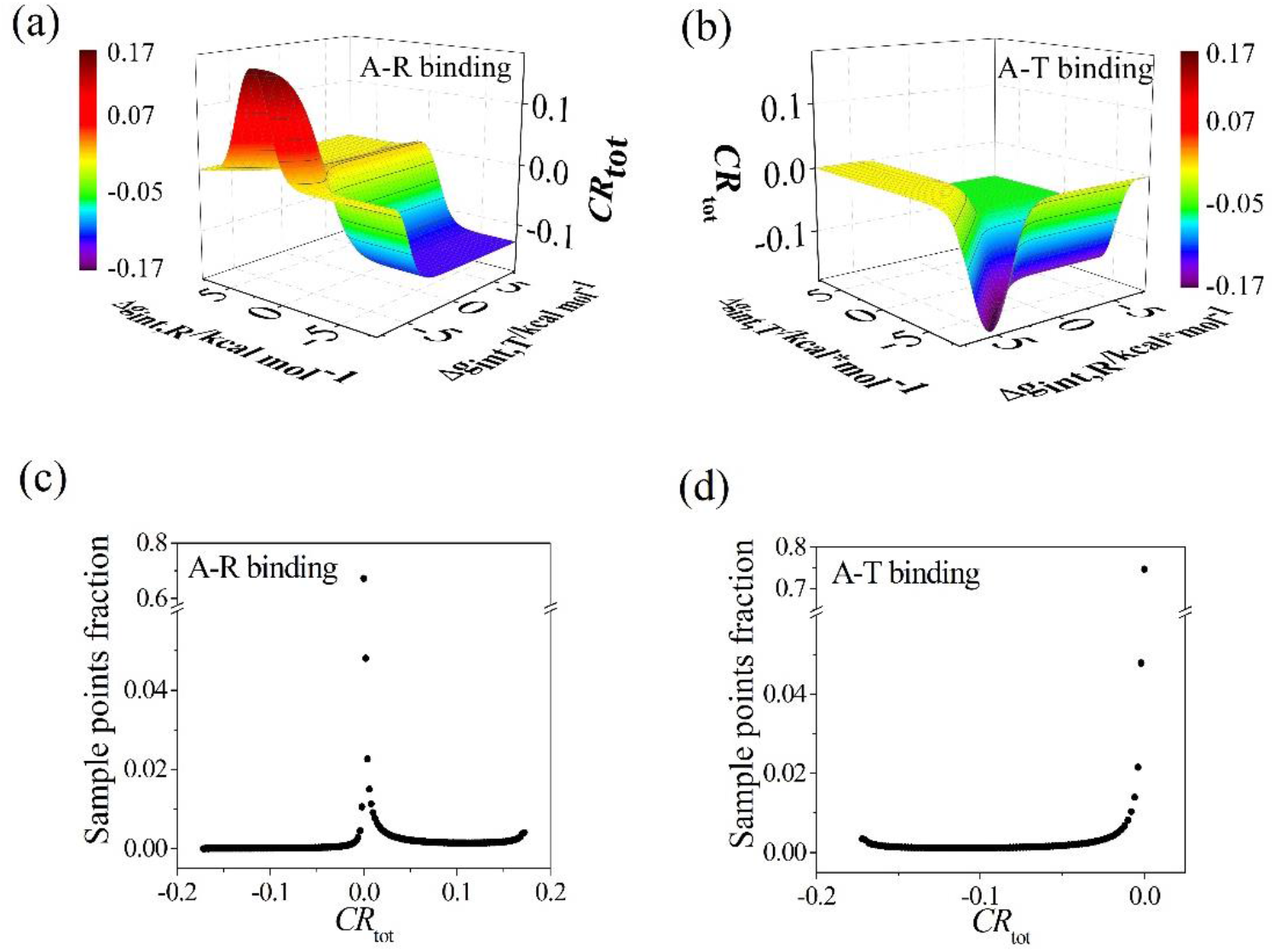
The allosteric coupling response (*CR*) of the comprehensive ensemble model. (a,b) *CR* as a function of Δ*g*_int,R_ and Δ*g*_int,T_ when the other parameters are fixed (chosen to produce notable allostery) as: Δ*G*_R1_ = −1.0, Δ*G*_R2_ = 1.3, Δ*G*_RT1_ = 1.0, Δ*G*_RT2_ = 3.0 (all in units of kcal/mol). Note that for A-T binding mode there is no activated allosteric effect. (c,d) Distribution of *CR* when the stability free-energy parameters (Δ*G*_R1_, Δ*G*_R2_, Δ*G*_RT1_, Δ*G*_RT2_, Δ*g*_int,R_, Δ*g*_int,T_) vary randomly with an equal probability density between −8 and +8 kcal/mol. The A-R binding mode is adopted in (a,c) and the A-T binding mode is adopted in (b,d) with Δ*g*_Lig,A_ = −3 kJ/mol.

The boundary limits of *CR* can be well explained in an analytic way. Take the MWC model as a simplified example, there are two states (RR and TT state) with only one stability parameter (Δ*G_i_*,. = G_*RR*_ – *G_TT_*), which determines the probability of RR state without ligand to be:

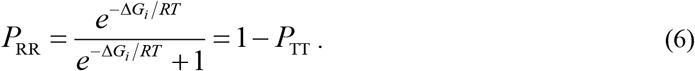

*CR* can then be written as a function of *P*_RR_ and Δ*g*_Lig,A_ as

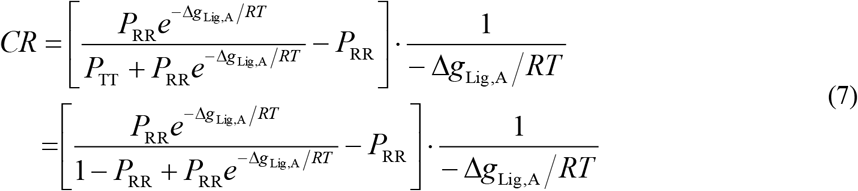

under the A-R binding mode. The relations among *P*_RR_, ΔG_i_ and *CR* are plotted in Fig. 5(a) for Δ*g*_Lig,A_ = −3 kJ/mol. *CR* is equal to 0 at either *P*_RR_ = 0 or *P*_RR_ = 1, i.e., too stable and too unstable RR state are unfavorable to allostery. *CR* reaches its maximum of about 0.172 at *P*_RR_ ≈ 0.081. *P*_RR_ depends on Δ*G_i_* in a switch-like manner. A great many Δ*G_i_* values give *P*_RR_ close to 0 or 1, and result in small *CR* and weak allostery. This provide a clue in understanding the dominant peak at *CR* = 0 in Fig. 4(c,d). Based on Eq. (7), the maximization of *CR* can be solved analytically with 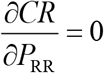 to give

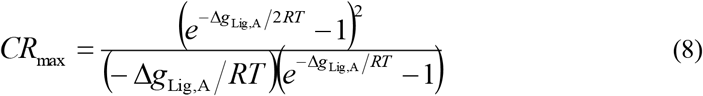

at the optimized *P*_RR_ as

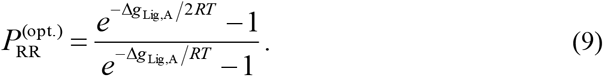

**Fig 5.**
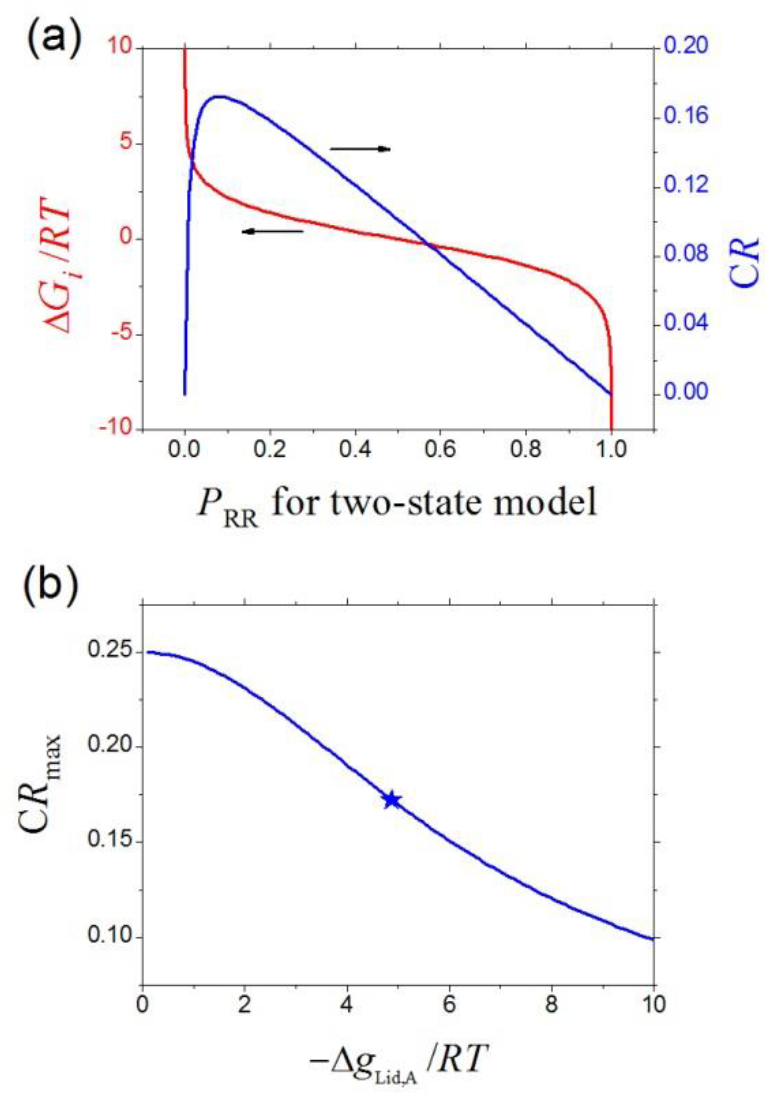
Limits of CR. (a) The relations among *P*_RR_, Δ*G_i_* and *CR* for the MWC model with two states, plotted according to Eqs. (6, 7). (b) The *CR* maximum (via optimizing state stabilities of proteins) as a function of −Δ*g*_Lig,A_/RT, plotted according to Eq. (8) which is valid for the comprehensive ensemble model as well as the MWC and EAM models. The star represents the data point for Δ*g*_Lig,A_ = −3 kJ/mol and *T* = 310.15 K used in other figures.

Eq. (8) keeps valid for the comprehensive ensemble model (see Supporting Information). *CR*_max_ is plotted in Fig. 5(b) as a function of −Δ*g*_Lig,A_/RT (note that Δ*g*_Lig,A_ < 0). It decreases with increasing −Δ*g*_Lig,A_/RT, and reaches a value of 0.172 at Δ*g*_Lig,A_ = −3 kJ/mol and *T* = 310.15 K, being consistent with the observation in Fig. 4. Eq. (8) gives an analytical result for the limits of *CR* when the state stabilities of protein are optimized, and would be useful in studying the allosteric capacity of proteins.

### The weight of MWC pathway is significantly higher than that of EAM pathway

The weights of three pathways (MWC, EAM and Others) in the allostery of the comprehensive system are numerically analyzed when the stability free-energy parameters (Δ*G*_R1_, Δ*G*_R2_, Δ*G*_RT1_, Δ*G*_RT2_, Δ*g*_int,R_, Δ*g*_int,T_) vary randomly between [−8, +8] kcal/mol. The resulting average weights are shown in Fig. 6(a) as functions of *CR*. For positive allosteric effect (*CR* > 0), the weight of the MWC pathway is much larger than the EAM one, indicating the MWC pathway holds an advantage over the EAM pathway in this case. For negative allosteric effect, *CR* under the A-R binding mode mainly comes from the EAM pathway, while under the A-T binding mode *CR* mainly comes from the MWC and Others pathways. The reason is that when A binds with R, in the MWC subsystem the decrease of RR state is not allowed and thus its weight is almost zero or even negative based on Eq. (5a), while an IR→RI transition of EAM pathway dominates the negative allosteric response. On the other hand, when A binds with T, it has no effect in the state distribution in the EAM subsystem thus its weight is always zero.

**Fig 6.**
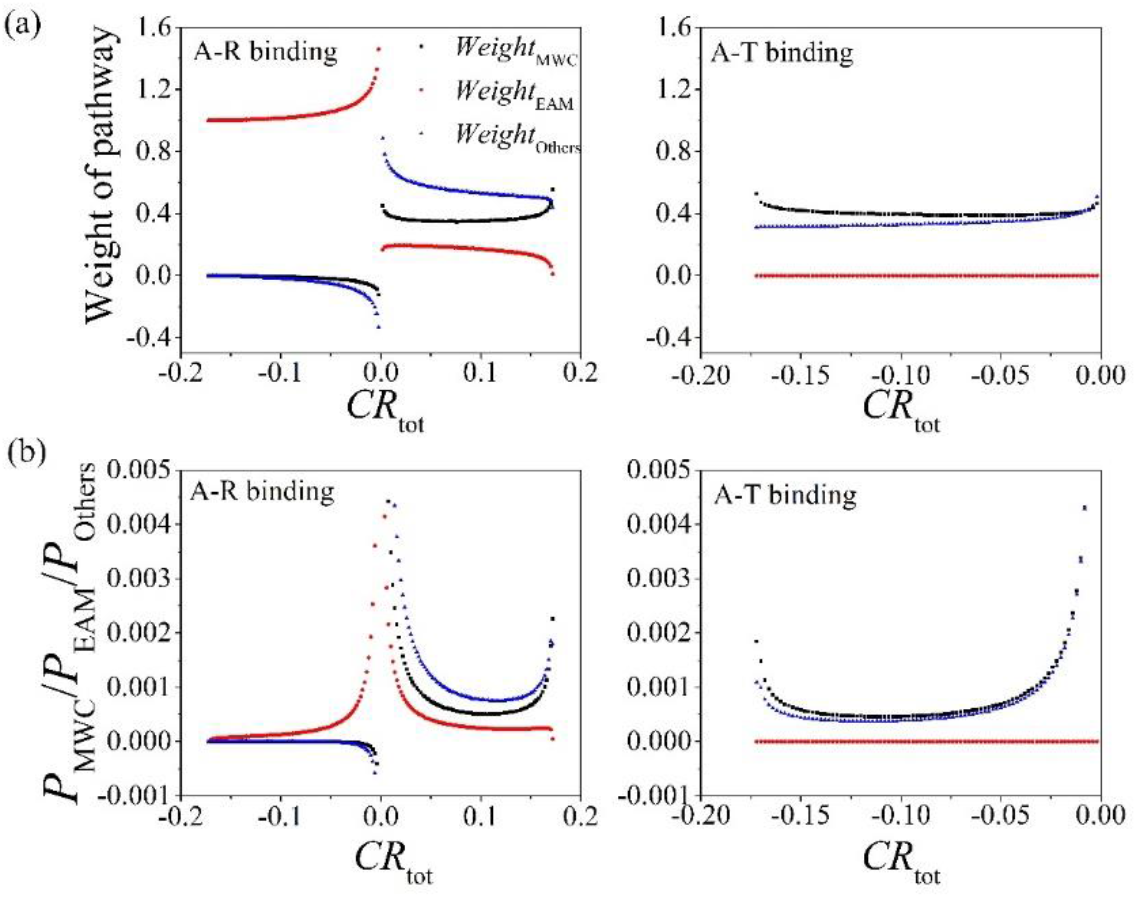
Contributions of three pathways (MWC, EAM, Others) in the comprehensive ensemble model. The stability free-energy parameters (Δ*G*_R1_, Δ*G*_R2_, Δ*G*_rT1_, Δ*G*_rT2_, Δ*g*_int,R_, Δ*g*_int,T_) vary randomly with an equal probability density between −8 and +8 kcal/mol, resulting in different systems (samples). (a,b) The average weights of pathways as functions of *CR*_tot_. The pathway weights of a system (sample) are calculated based on Eq. (5). (c,d) The capacity of three pathways for allostery, being calculated with Eq. (10). The A-R binding mode is adopted in (a,c) and the A-T binding mode is adopted in (b,d) with Δ*g*_Lig,A_ = −3 kJ/mol. It is noted that a large portion of samples practically have *CR* = 0 and the pathway contributions are ill-defined with Eq. (5), which are thus ignored.

The capacity of the MWC or the EAM pathway for allostery depends on not only their weights in a comprehensive system [Fig. 6(a)] but also the possibility of the system to afford an allosteric effect [*P*(*CR*), see Fig. 4(b)]. Therefore, the possibility for allosteric effect with *CR* undertaken by the MWC pathway can be calculated as

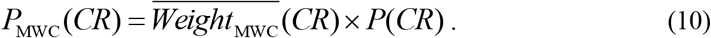

It describes the probability of a randomly chosen parameter set to possess an allosteric effect *CR* via the MWC pathway. Formula for the EAM and Others pathways can be similarly written. The calculated results are shown in Fig. 6(b). *P*_MWC_(*CR*) and *P*_Others_(*CR*) has sharp peak near the positive allostery limit *CR*_max_ in the A-R binding mode and near the negative allostery limit −*CR*_max_ in the A-T binding mode, which will be discussed in detail below. More importantly, if we take a simplified approach by adding curves in the A-R and A-T binding modes for each pathway, *P*_MWC_(*CR*) is much larger than *P*_EAM_(*CR*) for strong allosteric effects. Therefore, the MWC pathway is more important in allosteric effects than the EAM pathway based on the comprehensive ensemble model.

### Probability of strong allostery first increases and then decreases when the Δ*G_i_* range increases

The distribution of allostery and pathway contribution were investigated above when the free-energy parameters (Δ*G*_R1_, Δ*G*_R2_, Δ*G*_RT1_, Δ*G*_RT2_, Δ*g*_int,R_, Δ*g*_int,T_) of the comprehensive model vary randomly in a range of [−8, +8] kcal/mol. The results may change under a different range. In Fig. 7(a), the possibilities for an allosteric effect to occur with *CR* undertaken by three pathways are plotted under various variation range [−Δ*G*_max_, +Δ*G*_max_] of the free-energy parameters. The sharp peaks of *P*_MWC_ and *P*_Others_ near the positive allostery limit (*CR*_max_ = 0.172) observed previously are absent when the variation range (Δ*G*_max_) is small, e.g., Δ*G*_max_ = 1 kcal/mol. In Fig. 7(b), the probabilities of *CR* > 0.171 for three pathways are plotted as a function of Δ*G*_max_. It clearly shows that the MWC and the Others pathways have a similar tendency: it first equals to zero before a critical Δ*G*_max_ (which is smaller for the MWC pathway), then increases quickly, and finally decreases slowly.

**Fig 7.**
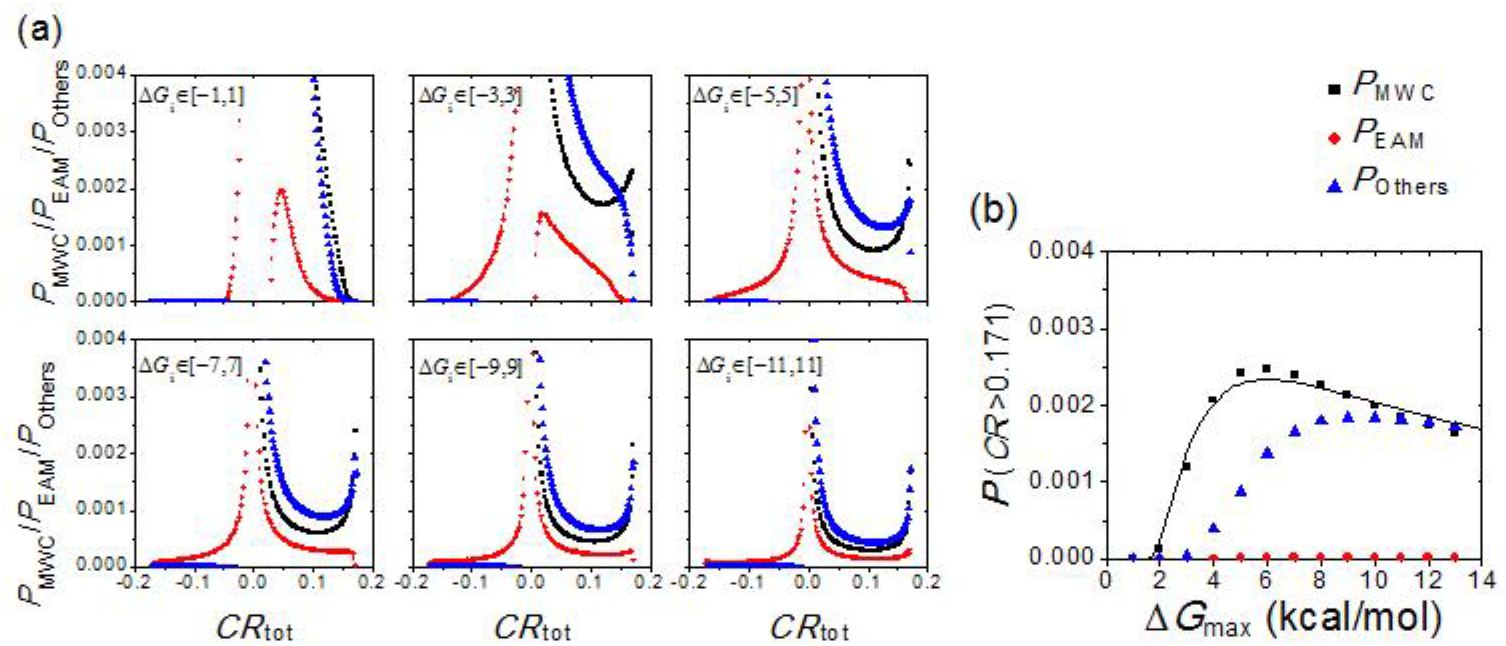
The influence of the variation range (Δ*G*_max_) for free-energy parameters (Δ*G*_R1_, Δ*G*_M_, Δ*G*_RT1_, Δ*G*_RT2_, Δ*g*_int,R_, Δ*g*_inr,T_) of the comprehensive ensemble model with the A-R binding mode. (a) The possibilities for three pathways [calculated with Eq. (10)] obtained at different [– Δ*G*_max_, +Δ*G*_max_] range. (b) The probabilities of *CR* > 0.171 as a function of ΔG_max_.

The feature observed in Fig. 7 can be qualitatively explained based on the simplified two-state model (Fig. 5). The maximal *CR* is achieved at *P*_RR_ = 0.081, which corresponds to a free energy difference of AG. (= *G*_RR_ – *G*_TT_) = 1.6 kcal/mol. When the variation range of the free-energy parameters is small, the resulting Δ*G_i_*, cannot reach the optimized value for the maximal *CR*, giving the zero value in Fig. 7(b) and the absence of the sharp peak near *CR*_max_ in the panel with Δ*G*_max_ = 1 kcal/mol in Fig. 7(a). When the variation range of the free-energy parameters is large enough, although the optimized value of Δ*G_i_*, can be always satisfied at some values of parameter sets, the total number of possible values increases with the variation range, and thus the probability of maximal *CR*, defined as the ratio between the number of optimized parameter value sets to that of the total number, would decreases with increasing the variation range as observed in Fig. 7(b).

### Two-state transition is the main mechanism for strong allostery

The comprehensive ensemble model includes seven states and three subsystems/pathways. How do they coordinate in fulfilling the allosteric effect? For example, do the pathways repeal each other in a system? How many states play significant role in a system? Here, we investigate the interplay between different states and different subsystems/pathways in the allosteric process.

To measure the mixing extend of subsystems and pathways, we classify each system case (with a certain set of Δ*G_i_* values) into one of four categories: single subsystem with single pathway (S,S), single subsystem with mixing pathways (S,M), mixing subsystems with single pathway (M,S), and mixing subsystems with mixing pathways (M,M). If the sum of state probability for any subsystem is larger than 0.99 before and after adding ligand, it is classified into single subsystem; otherwise it belongs to mixing subsystem. Single pathway is defined for the case where the weight of one pathway is larger than 0.99 and the absolute value of weights for other pathways are less than 0.01; otherwise it belongs to mixing pathway. For example, if a system only contain RR and TT states, then it simply belongs to the (S,S) category. The results are shows in Fig. 8(a). When the variation range (Δ*G*_max_) of free-energy parameters is small, mixing subsystems with mixing pathways (M,M) dominate in most cases. But when Δ*G*_max_ is larger, the proportion of single subsystem with single pathway (S,S) increases while the (M,M) type decreases. More importantly, the (S,S) proportion increases with increasing |*CR*|. The system tends to behave as pure subsystem with pure pathway mechanism at strong allostery.

**Fig 8.**
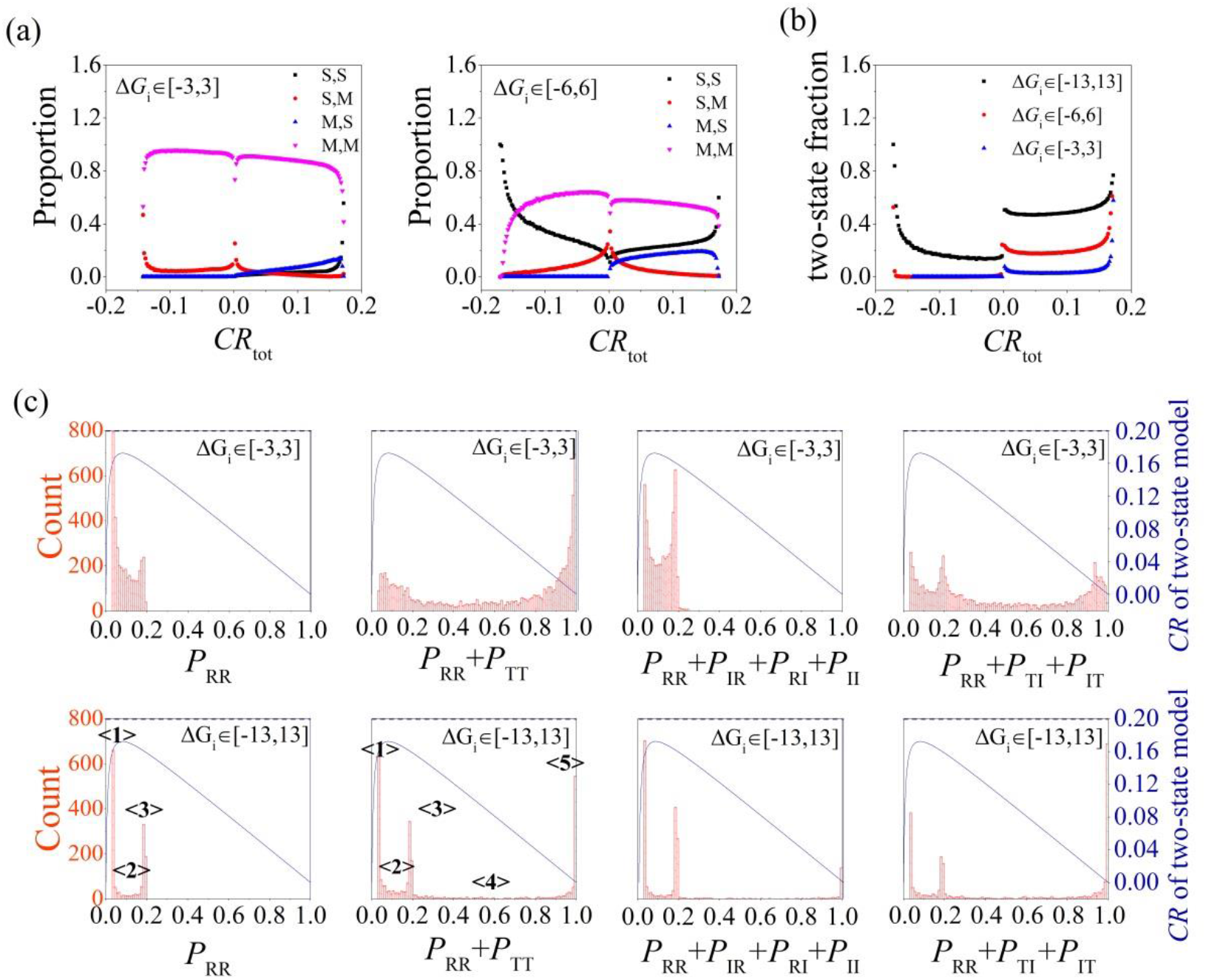
Interplay between different states and subsystems/pathways in the comprehensive ensemble model. The A-R binding mode is adopted. (a) Proportion of four categories [single subsystem with single pathways (S,S), single subsystem with mixing pathways (S,M), mixing subsystems with single pathways (M,S), and mixing subsystems with mixing pathways (M,M)] in systems with *CR*. (b) The proportion of systems with two-state transition. (c) Distribution of *P*_RR_ and state probability of three subsystems (*P*_RR_ + *P*_TT_ for the MWC pathway, *P*_RR_ + *P*_IR_ + *P*_RI_ + *P*_II_ for the EAM pathway, and *P*_RR_ + *P*_TI_ + *P*_IT_ for the Other pathway) for systems with *CR* ≈ 0. 16. The theoretical *CR* ~ *P*_RR_ curve (blue line) for the two-state model is also plotted using Eq. (7). The horizontal dashed line represents *CR* = 0.16.

A clearer angle of view is to look at the proportion of systems that implement allostery via a simple mechanism of two-state transition. Here we specify a system to have two-state transition mechanism if the probability sum of two certain states of the given system is larger than 0.99 both before and after binding with ligand A. Possible two-state transition for positive allosteric effect includes “II➔RR”, “TT➔RR”, “TI➔RR” and “IT➔RR” For negative allosteric effect, the only possible two-state transition is “IR➔RI”. The proportion of systems with simple two-state transition is shown in Fig. 8(b). With larger Δ*G*_max_, the proportion of two-state transition is higher. The proportion has a sharp peak at ±*CR*_max_. Therefore, two-state transition is the major mechanism for strong allosteric even in the comprehensive ensemble model.

The existence of two-state transition and single subsystem/pathway are also reflected in the state distribution patterns. The distributions of RR and states of three subsystems are shown in Fig. 8(c) for systems with *CR* ≈ 0.16. The distribution of *P*_RR_ has two obvious peaks labeled with <**1**> and <**3**>. In Fig. 8(c) we also plot the theoretical *CR* ~ *P*_RR_ curve for the two-state model for convenience’s sake. The crossing points between the *CR* ~ *P*_RR_ curve and the horizontal line of *CR* = 0.16 give the P_RR_ values to achieve an allosteric effect of *CR* = 0.16 in the two-state model. The obtained *P*_RR_ values of the crossing points coincide with the peak position at <**1**> and <**3**> of the simulated *P*_RR_ distribution, suggesting that the strong allostery (with *CR* = 0.16) of the comprehensive model mainly occurs in a two-state model mechanism (note that RR exists in all possible two-state transition for positive *CR* including “II➔RR”, “TT➔RR”,”TI➔RR” and “IT➔RR”). There is also some nonzero *P*_RR_ distribution (<**2**>) between two peaks, which is expected to have *CR* higher than 0.17 in the two-state model. The reason for that is the introducing of additional IR and RI population would decrease *CR* (see Supplementary Material). It also explains the intriguing result that there is no distribution outside <**1**>&<**3**>, for *P*_RR_ outside cannot give *CR* as big as 0.16. When Δ*G*_max_ increases to 13 kcal/mol, P_RR_ distribution enriches at <**1**>&<**3**> and reduces at <**2**>, suggesting an enrichment of two-state transition mechanism. Similarly, for the distribution of the MWC pathway states, the *P*_RR_ + *P*_TT_ peaks at <**1**>&<3> correspond to the systems dominated by other pathways (EAM or Others) so that *P*_RR_ + *P*_TT_ = *P*_RR_ and the peak positions are identical to that for *P*_RR_. At <5>, *P*_RR_ + *P*_TT_ = 1 corresponds to the systems dominated by the MWC pathway. <**2**> and <**4**> mean hybridized cases. Results for the population distribution of the EAM and Others subsystems are similar (data not shown). They confirm that strong allostery in the comprehensive ensemble mode is dominated by single pathway and the two-state transition mechanism.

## Discussion

### Possible reasons for the prevalence of IDPs/IDRs in allosteric regulation

IDPs/IDRs appear in much higher amounts in regulatory proteins,(20, 23) and are also widely involved in allosteric processes.(35–40, 42) A possible explain for the prevalence of IDPs/IDRs in allosteric regulation was provided by the EAM model which suggested that intrinsic disorder can maximize the ability to allosteric coupling.(45) However, our comprehensive ensemble model reveals that the order-disorder transition (EAM mechanism) is actually less competitive than the order-order transition (MWC mechanism) in affording allosteric effects, especially the strong allostery. It shows that the reasons for the prevalence of IDPs/IDRs in allosteric regulation are more complicated than previously thought. Our work does not give a complete answer for it, but we provide some discussion and comments here.

Firstly, in our study we assumed that the free energy parameters of conformation change and domain-domain interaction (Δ*G*_R1_, Δ*G*_R2_, Δ*G*_RT1_, Δ*G*_RT2_, Δ*g*_int,R_, Δ*g*_int,T_) vary randomly with an equal probability density between [−Δ*G*_max_, +Δ*G*_max_]. In real proteins it does not have to be like this. The difficulty (probability) to modify order-order and order-disorder transitions is likely different. Specifically, to tune the protein stability difference between two similar order structures (R and T in the MWC model) via mutation would be more difficult than to tune the stability difference between order and disordered structures (R and I in the EAM model), because in the latter case this can be accomplished via breaking or strengthening a residue-residue interaction that is present in ordered structure but absent in disordered structure. Therefore, a possible reason for the prevalence of IDPs/IDRs in allosteric regulation is their convenience in modifying state stability.

Secondly, IDPs/IDRs possess various advantages over ordered proteins,(24, 50) such as saving genome resources via multi-binding pattern or creating large binding surface, overcoming steric effect in binding, accelerating binding speed, achieving high specificity with low affinity, and facilitating posttranslational modifications. The prevalence of IDPs/IDRs in allosteric regulation is determined by all their advantage, but not only by their capacity in endowing allostery. Work combining experimental data and bioinformatics analyses would be helpful to compare ordered and disordered proteins’ importance in allosteric regulation.

Lastly, allosteric effects with maximal *CR* may be not the pursuing goal. Allostery with different strength would have different applications. For example, allosteric effect that are not too strong is beneficial in ensuring safer dosing.(51)

## Conclusions

In this work, we proposed a comprehensive ensemble model to study the role of order-order and order-disorder transitions in allosteric effect. An analytic equation for the maximal allosteric coupling response (*CR*) was derived, which shows that too stable or too unstable state is unfavorable to achieve allostery. By sampling the parameter space, it was revealed that the order-order transition (MWC) mechanism has a higher possibility in allostery than the order-disorder transition (EAM) mechanism. In addition, two-state transition is the primary mechanism when allostery is strong although there are seven states in the model. The work not only provided insight in understand the prevalence of IDPs/IDRs in allosteric regulation, but is also helpful for rational design of allosteric drugs.

## Acknowledgments

This work was supported by the National Natural Science Foundation of China (grant 21633001) and the Ministry of Science and Technology of China (grant 2015CB910300). The authors thank Huaiqing Cao, Miao Yu and Hao Ruan for helpful discussions.

## Supporting Information figure captions

Fig. S1 CR Distribution of CRMWC and CREAM for the separated MWC and EAM subsystems. Parameters are identical to those in Fig. 4.

## References

1. Fenton AW. Allostery: an illustrated definition for the ‘second secret of life’. Trends Biochem Sci. 2008;33(9):420–5.

2. Monod J, Changeux JP, Jacob F. Allosteric proteins and cellular control systems. J Mol Biol. 1963;6(4):306–29.

3. Monod J, Wyman J, Changeux JP. On nature of allosteric transitions - a plausible model. J Mol Biol. 1965;12(1):88–118.

4. Koshland DE, Nemethy G, Filmer D. Comparison of experimental binding data and theoretical models in proteins containing subunits. Biochemistry. 1966;5(1):365–85.

5. Eaton WA, Henry ER, Hofrichter J, Mozzarelli A. Is cooperative oxygen binding by hemoglobin really understood? Nat Struct Biol. 1999;6(4):351–8.

6. Iwata S, Kamata K, Yoshida S, Minowa T, Ohta T. T-state and R-state in the crystals of bacterial l-lactate dehydrogenase reveal the mechanism for allosteric control. Nat Struct Biol. 1994;1(3):176–85.

7. Lockless SW, Ranganathan R. Evolutionarily conserved pathways of energetic connectivity in protein families. Science. 1999;286(5438):295–9.

8. Tang S, Liao J-C, Dunn AR, Altman RB, Spudich JA, Schmidt JP. Predicting allosteric communication in myosin via a pathway of conserved residues. J Mol Biol. 2007;373(5):1361–73.

9. Bochkareva E, Belegu V, Korolev S, Bochkarev A. Structure of the major single-stranded DNA-binding domain of replication protein A suggests a dynamic mechanism for DNA binding. EMBO J. 2001;20(3):612–8.

10. Lukin JA, Kontaxis G, Simplaceanu V, Yuan Y, Bax A, Ho C. Quaternary structure of hemoglobin in solution. Proc Natl Acad Sci. 2003;100(2):517–20.

11. Tsai C-J, Nussinov R. A unified view of “how allostery works”. PLoS Comput Biol. 2014;10(2):e1003394.

12. Wright PE, Dyson HJ. Intrinsically unstructured proteins: Re-assessing the protein structure-function paradigm. J Mol Biol. 1999;293(2):321–31.

13. Dunker AK, Lawson JD, Brown CJ, Williams RM, Romero P, Oh JS, et al. Intrinsically disordered protein. J Mol Graph Model. 2001;19(1):26–59.

14. Huang Y-Q, Liu Z-R. Intrinsically disordered proteins: the new sequence-structure-function relations. Acta Phys-Chim Sin. 2010;26(8):2061–72.

15. Tompa P. Unstructural biology coming of age. Curr Opin Struc Biol. 2011;21(3):419–25.

16. Huang Y, Yoon M-K, Otieno S, Lelli M, Kriwacki RW. The activity and stability of the intrinsically disordered Cip/Kip protein family are regulated by non-receptor tyrosine kinases. J Mol Biol. 2015;427(2):371–86.

17. Wallin S. Intrinsically disordered proteins: structural and functional dynamics. Res Rep Biol. 2017;8:7–16.

18. Csizmok V, Follis AV, Kriwacki RW, Forman-Kay JD. Dynamic protein interaction networks and new structural paradigms in signaling. Chem Rev. 2016;116(11):6424–62.

19. Uversky VN. Natively unfolded proteins: A point where biology waits for physics. Protein Sci. 2002;11(4):739–56.

20. Liu J, Perumal NB, Oldfield CJ, Su EW, Uversky VN, Dunker AK. Intrinsic disorder in transcription factors. Biochemistry. 2006;45(22):6873–88.

21. Uversky VN, Oldfield CJ, Dunker AK. Showing your ID: intrinsic disorder as an ID for recognition, regulation and cell signaling. J Mol Recognit. 2005;18.

22. Uversky VN. A decade and a half of protein intrinsic disorder: Biology still waits for physics. Protein Sci. 2013;22(6):693–724.

23. Xie H, Vucetic S, Iakoucheva LM, Oldfield CJ, Dunker AK, Uversky VN, et al. Functional anthology of intrinsic disorder. 1. Biological processes and functions of proteins with long disordered regions. J Proteome Res. 2007;6(5):1882–98.

24. Liu Z, Huang Y. Advantages of proteins being disordered. Protein Sci. 2014;23(5):539–50.

25. Huang Y, Liu Z. Do intrinsically disordered proteins possess high specificity in protein-protein interactions? Chemistry. 2013;19(14):4462–7.

26. Zhou H-X. Intrinsic disorder: signaling via highly specific but short-lived association. Trends Biochem Sci. 2012;37(2):43–8.

27. Wong ETC, Na D, Gsponer J. On the importance of polar interactions for complexes containing intrinsically disordered proteins. PLoS Comput Biol. 2013;9(8):e1003192.

28. Bhattacherjee A, Wallin S. Exploring protein-peptide binding specificity through computational peptide screening. PLoS Comput Biol. 2013;9(10): e1003277.

29. Karush F. Heterogeneity of the binding sites of bovine serum albumin. J Am Chem Soc. 1950;72(6):2705–13.

30. Kriwacki RW, Hengst L, Tennant L, Reed SI, Wright PE. Structural studies of p21^(Waf1/Cıp1/Sdl1)^ in the free and Cdk2-bound state: Conformational disorder mediates binding diversity. Proc Natl Acad Sci. 1996;93(21):11504–9.

31. Borg M, Mittag T, Pawson T, Tyers M, Forman-Kay JD, Chan HS. Polyelectrostatic interactions of disordered ligands suggest a physical basis for ultrasensitivity. Proc Natl Acad Sci. 2007;104(23):9650–5.

32. Cortese MS, Uversky VN, Dunker AK. Intrinsic disorder in scaffold proteins: Getting more from less. Prog Biophys Mol Bio. 2008;98(1):85–106.

33. Tompa P, Csermely P. The role of structural disorder in the function of RNA and protein chaperones. FASEB J. 2004;18(11):1169–75.

34. Xue B, Dunker AK, Uversky VN. The roles of intrinsic disorder in orchestrating the Wnt-pathway. J Biomol Struct Dyn. 2012;29(5):843–61.

35. Tompa P. Multisteric regulation by structural disorder in modular signaling proteins: an extension of the concept of allostery. Chem Rev. 2014;114(13):6715–32.

36. Reichheld SE, Yu Z, Davidson AR. The induction of folding cooperativity by ligand binding drives the allosteric response of tetracycline repressor. Proc Natl Acad Sci. 2009;106(52):22263–8.

37. Freiburger LA, Baettig OM, Sprules T, Berghuis AM, Auclair K, Mittermaier AK. Competing allosteric mechanisms modulate substrate binding in a dimeric enzyme. Nat Struct Mol Biol. 2011;18(3):288–94.

38. Garcia-Pino A, Balasubramanian S, Wyns L, Gazit E, De Greve H, Magnuson RD, et al. Allostery and intrinsic disorder mediate transcription regulation by conditional cooperativity. Cell. 2010;142(1):101–11.

39. Smock RG, Gierasch LM. Sending Signals Dynamically. Science. 2009;324(5924):198–203.

40. Krishnan N, Koveal D, Miller DH, Xue B, Akshinthala SD, Kragelj J, et al. Targeting the disordered C terminus of PTP1B with an allosteric inhibitor. Nat Chem Biol. 2014;10(7):558–66.

41. Liu J, Nussinov R. Allosteric effects in the marginally stable von Hippel-Lindau tumor suppressor protein and allostery-based rescue mutant design. Proc Natl Acad Sci. 2008;105(3):901–6.

42. Garcia-Pino A, De Gieter S, Talavera A, De Greve H, Efremov RG, Loris R. An intrinsically disordered entropic switch determines allostery in Phd-Doc regulation. Nat Chem Biol. 2016;12(7):490–6.

43. Kern D, Zuiderweg ERP. The role of dynamics in allosteric regulation. Curr Opin Struc Biol. 2003;13(6):748–57.

44. Gunasekaran K, Ma BY, Nussinov R. Is allostery an intrinsic property of all dynamic proteins? Proteins. 2004;57(3):433–43.

45. Hilser VJ, Thompson EB. Intrinsic disorder as a mechanism to optimize allosteric coupling in proteins. Proc Natl Acad Sci. 2007;104(20):8311–5.

46. Motlagh HN, Wrabl JO, Li J, Hilser VJ. The ensemble nature of allostery. Nature. 2014;508(7496):331–9.

47. Choi JH, Laurent AH, Hilser VJ, Ostermeier M. Design of protein switches based on an ensemble model of allostery. Nat Commun. 2015;6:1–9.

48. Motlagh HN, Li J, Thompson EB, Hilser VJ. Interplay between allostery and intrinsic disorder in an ensemble. Biochem Soc Trans. 2012;40:975–80.

49. Maillard RA, Liu T, Beasley DWC, Barrett ADT, Hilser VJ, Lee JC. Thermodynamic mechanism for the evasion of antibody neutralization in flaviviruses. J Am Chem Soc. 2014;136(29):10315–24.

50. Huang Y, Liu Z. Kinetic advantage of intrinsically disordered proteins in coupled folding-binding process: A critical assessment of the “fly-casting” mechanism. J Mol Biol. 2009;393(5):1143–59.

51. Wagner JR, Lee CT, Durrant JD, Malmstrom RD, Feher VA, Amaro RE. Emerging computational methods for the rational discovery of allosteric drugs. Chem Rev. 2016;116(11):6370–90.

